# Prioritizing landscapes to reconcile biodiversity conservation, ecosystem services, and human well-being in India

**DOI:** 10.1101/2022.08.27.505513

**Authors:** Arjun Srivathsa, Divya Vasudev, Tanaya Nair, Stotra Chakrabarti, Pranav Chanchani, Ruth DeFries, Arpit Deomurari, Sutirtha Dutta, Dipankar Ghose, Varun R. Goswami, Rajat Nayak, Amrita Neelakantan, Prachi Thatte, Srinivas Vaidyanathan, Madhu Verma, Jagdish Krishnaswamy, Mahesh Sankaran, Uma Ramakrishnan

**Affiliations:** National Centre for Biological Science, TIFR, Bengaluru, India; Conservation Initiatives, Guwahati, India; Division of Biosciences, University College London, London, UK; Departments of Biology & Environmental Studies, Macalester College, Minnesota, USA; World Wildlife Fund, Delhi, India; Department of Ecology, Evolution, and Environmental Biology, Columbia University, New York, USA; RoundGlass H2O Pvt. Ltd, Mohali, India; Wildlife Institute of India, Dehradun, India; Foundation for Ecological Research, Advocacy and Learning, Bengaluru, India; Network for Conserving Central India, India; World Resources Institute, New Delhi, India; School of Environment and Sustainability, Indian Institute for Human Settlements, Bengaluru, India; Ashoka Trust for Research in Ecology and the Environment, Bengaluru, India; Biodiversity Collaborative, India

**Keywords:** ecosystem services, global biodiversity targets, landscape conservation, spatial prioritization, sustainable development

## Abstract

Biodiversity conservation and human well-being are tightly interlinked; yet mismatches in the scale at which both priorities are planned and implemented have exacerbated biodiversity loss, erosion of ecosystem services, and declining human quality of life. India houses the second largest human population on the planet, while <5% of the country’s land area is effectively protected for conservation. This warrants landscape-level conservation planning through a judicious mix of *land-sharing* and *land-sparing* approaches, and co-production of ecosystem services. Through a multi-faceted assessment, we prioritize spatial extents of land parcels that, in the face of anthropogenic threats, can safeguard conservation landscapes across India’s biogeographic zones. We find that only a fraction (~15%) of such priority areas identified here are encompassed under India’s extant PA network, and several landscapes of high importance were omitted in all previous global-scale assessments. We then examined the spatial congruence of priority areas with administrative units earmarked for economic development by the Indian government, and propose management-zoning through state-driven and participatory approaches. Our spatially explicit insights can help meet the twin goals of biodiversity conservation and sustainable development in India and other countries across the Global South.

## Main

Concomitant impacts of biodiversity decline, climate change, unsustainable land use, and inequitable extraction of natural resources, have degraded the quality of human life (Levett 2002; Harrison 2010; Otero et al. 2020). Biodiversity underpins several provisioning, regulatory, and cultural or aesthetic ecosystem services. Ecosystem services linked to biodiversity, in turn, are crucial for ensuring long-term human well-being (Seppelt et al. 2013). Recent discourse on nature-based solutions acknowledges nature’s contributions as humanity’s ‘safety net’ (IPBES 2019; Dinerstein et al. 2020), and that ecological processes are shaped by a complex interplay between ecological *and* social systems (Ostrom 2009; Summers et al. 2012; Bennett et al. 2015). A sustainable future can only be ensured by adopting an ecosystems approach to conservation that emphasizes links between human and natural systems to address global, regional and biome-scale threats to their functioning (Haines-Young & Potschin 2010). Siloed management approaches to meet conservation targets that ignore this interplay have failed to yield equitable and sustainable progress (Tallis & Kareiva 2006; Steffen et al. 2018). A renewed global resolve will need to consider nature in the Anthropocene, and explicitly reconcile land-use planning for economic development with ecosystem functioning, biodiversity conservation and long-term human well-being.

For nearly two decades, several priority-setting tools have been proposed the world over to identify and rank species, habitats and locations based on their relative importance, vulnerability, or ease of management interventions (Wilson et al. 2007). These approaches generally adopt systematic spatial conservation planning––a set of decision-making tools that help determine the spatial locations where resources and actions need to be directed to optimize conservation gains (Moilanen et al. 2005, 2011; Game et al. 2013). Earlier iterations of these tools largely involved identifying locations to demarcate Protected Areas (PAs), determine optimal reserve design, or propose conservation corridors (Kremen et al. 2008). But human interests like economic aspirations, infrastructure development, and agricultural expansion to ensure food security must be integrated with conservation goals to enhance traction with policy makers (Pressey et al. 2007). This necessitates adopting *landscape approaches* to conservation (see Sayer et al. 2013), so as to offer pragmatic solutions in the Anthropocene. Such considerations led to the idea of ‘zoning’ landscapes, whereby assessors not only identify priority locations, but also stratify them into various zones and determine appropriate management interventions (Watts et al. 2009; Arkema et al. 2015). This combination of prioritization–zoning therefore holds promise to guide better management of landscapes in an increasingly human-modified world.

Since its conception, the idea of prioritizing areas for conservation at the global level has seen wide application in scientific literature. While these investigations can offer important insights at macro-scales, their real-world applications have been limited (Wyborn & Evans 2021). One reason for this is perhaps the lack of alignment between scientific ‘boundaries’ (sampling units) and administrative jurisdictions; yet, it is within these jurisdictions that implementation typically occurs (Seppelt et al. 2013). Aligning ecological findings with administrative boundaries, or existing policy, may thus increase the utility of prioritization exercises. Second, making assessments at global scales may compromise the spatial resolution of analyses, hampered by the availability and comparability of data at scale, or miss out on local socio-political nuances (Brooks et al. 2006; Turner et al. 2007; Jenkins et al. 2013; Brum et al. 2017). Third, macro-scale analyses could entail intrinsic biases in representation of key features or attributes. For example, certain historically overlooked biomes may continue to remain ignored (Silviera et al. 2021); habitats presumed to be ‘unproductive’ may not be prioritized (e.g., grasslands, see Bond & Parr 2010); indices such as species richness, which ignore community composition, may discount rarity or endemism (Veach et al. 2017); PAs or ‘intact’ wilderness areas may take precedence over heterogeneous multi-use conservation landscapes (Venter et al. 2014; Potapov et al. 2017); and important dimensions such as human populations may altogether be excluded (Jung et al. 2021).

Achieving conservation goals, sustaining ecosystem services, and ensuring human well-being while balancing economic development present formidable challenges in implementation (Otero et al. 2020). India exemplifies this premise; (i) it is a large, diverse country with ten distinct biogeographic zones and four biodiversity hotspots; (ii) its Protected Area (PA) network, conservation landscapes and riverscapes support several ecologically important and evolutionarily distinct species assemblages; (iii) it has the second largest human population in the world, with a large proportion of the people directly dependent on natural resources drawn from natural ecosystems; and (iv) with rapid infrastructure development and liberal investment policies, India is currently among the fastest-growing economies in Asia. In this study, we: (1) identify spatial scale(s), resolutions, and thematic dimensions for a priority-setting exercise across the country (Fig. 1); (2) use a systematic spatial prioritization approach to optimize landscapes for habitat protection, biodiversity conservation, and ecosystem service gains, while penalizing locations facing negative anthropogenic impacts; and (3) align our results with global biodiversity targets (Post-2020 Global Biodiversity Framework, CBD 2020) and the Indian government’s development initiative to identify ‘aspirational’ economic districts. We provide insights that can guide future environmental planning and policy to help meet the twin goals of biodiversity conservation and sustainable development in a strategically important country for Asia. More broadly, our framework can be adapted and applied for conservation planning in other eco-regions of developing countries.

**Figure 1.**
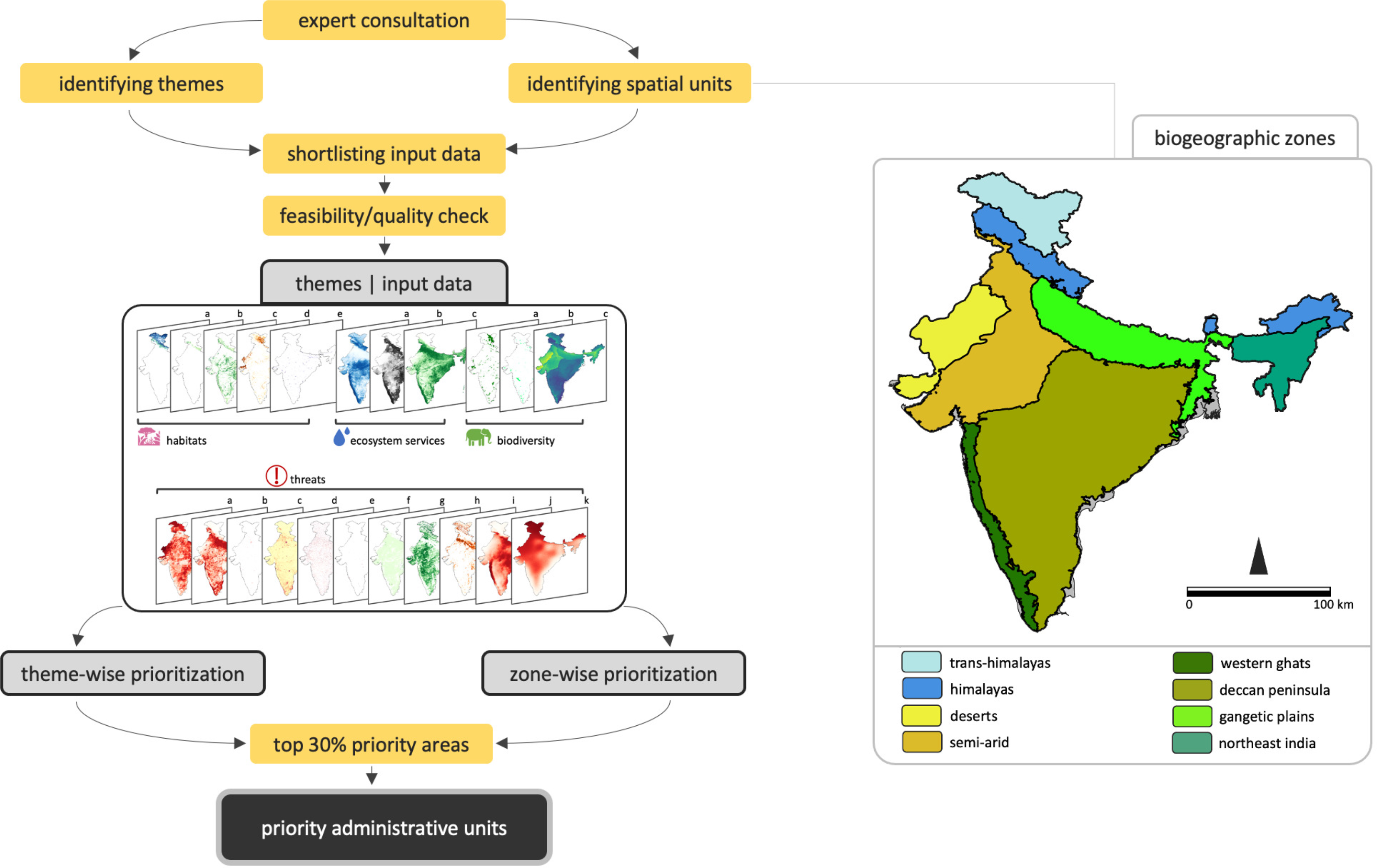
Schematic of the study framework, biogeographic zones of India, and themes and input layers used for spatial prioritization. Theme-wise input layers include: (1) Habitats: 28 vegetation classes depicted as 5 composite panels; (2) Ecosystem Services: blue water flux, above- and below-ground carbon, and green water flux; (3) Biodiversity: Protected Areas, Key Biodiversity Areas, and a diversity index of threatened mammals, birds, reptiles and amphibians; (4) Threats: human population density, livestock population density, urbanization, linear infrastructure, mines, river fragmentation, agricultural expansion, vegetation greening, vegetation browning, future climate warming and future rainfall anomaly.

## Results and Discussion

### a. Overlap in priority sites across themes

Priority maps generated independently for habitats, ecosystem services and biodiversity (Fig. 2a) represented sites of high value in isolation, i.e., sites that were: (a) representative of important and rare natural habitats, (b) responsible for the provision of key ecosystem services, viz. water and carbon, and (c) important solely from the perspective of threatened species diversity and turnover. We found moderate overlap amongst these three layers, which varied across biogeographic zones (hereafter, ‘zones’; Fig. 2). The overlap of priority sites for biodiversity (defined as the top 30% rankings) and habitats was 37% across the country, ranging from 30% in deserts, to 48% in the Western Ghats (Fig. S1 in Supplementary File 1). There was significant addition of sites when ecosystem services was added as a criterion, in addition to biodiversity and habitats. Around 38% of priority sites for ecosystem services were not covered by either biodiversity or habitat priority sites; these locations collectively contain a population of 2.3 million people (based on spatial overlap alone) who are directly dependent on these watersheds (see Fig. S1).

**Figure 2.**
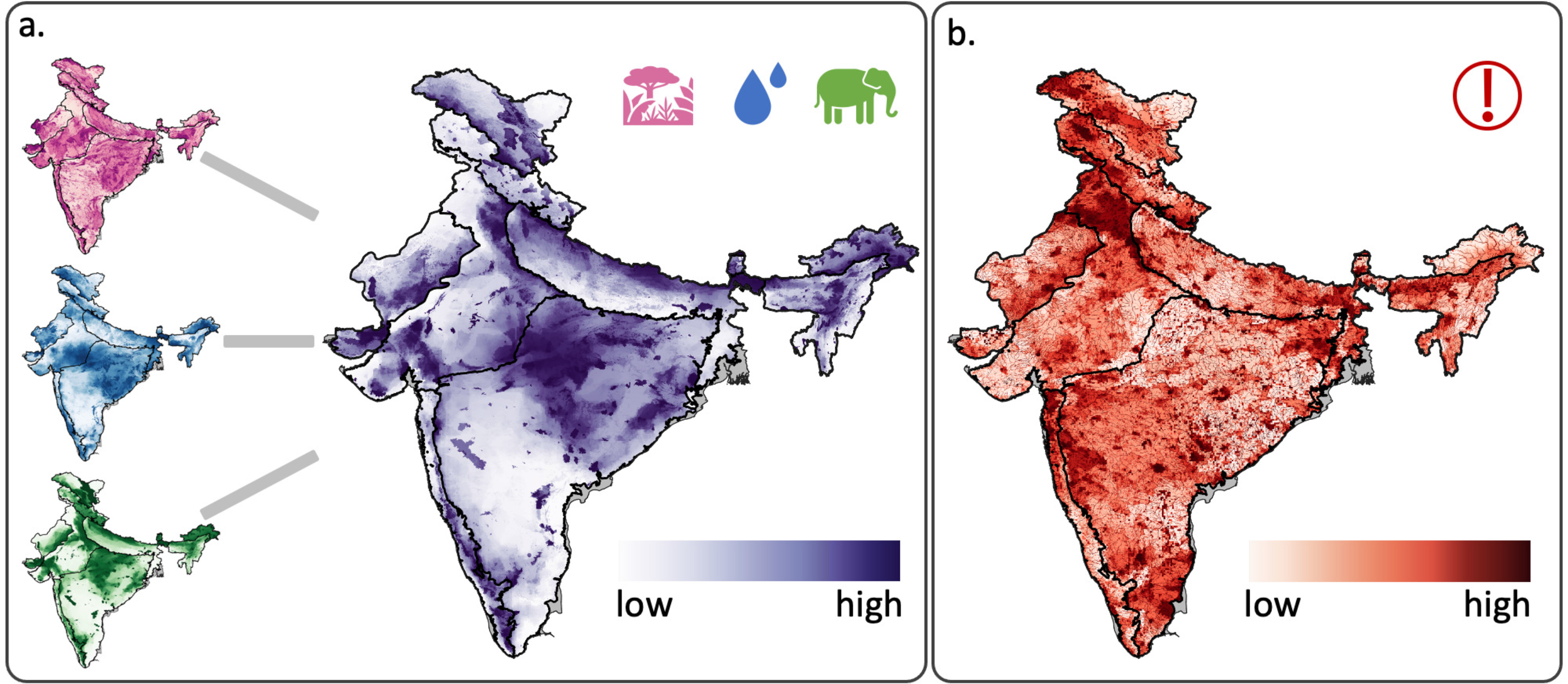
(a) Theme-wise results from prioritization analyses of Habitats (top left), Ecosystem services (centre left), Biodiversity (bottom left), and Composite of the three themes (right); (b) Theme-specific results from prioritization analysis of Threats.

Conservationists often highlight the mismatches in priority sites when different criteria are used to assign a ‘conservation value’ (Westgate et al. 2014; Cadotte & Tucker 2018). Our findings support this concern (Fig. 2) and point to the value of incorporating multiple contributors to conservation value. Of significance, our multi-criterion approach selected for India’s Open Natural Ecosystems (Madhusudan & Vanak 2021) — prioritized based on habitats, but not based on biodiversity — encompassing open grasslands, savannas, hot and cold deserts, ravines, rocky boulders and escarpments. These systems are extremely fragile, with unique endemic flora and fauna, but inappropriately classified as “wastelands” as per India’s land-use and conservation policy (Wastelands Atlas of India 2019). Similarly, areas ranked high for habitat and biodiversity, but low in terms of ecosystem services represented some locations in the arid/semi-arid dry zones of western India, the cold deserts of trans-Himalayas, and parts of the terai grasslands along the India-Nepal border (Fig. 2a). Conversely, areas ranked high for ecosystem services, critical for water and carbon storage, did not rank high for biodiversity or habitats in some places (e.g., parts of Northeast India). This, perhaps, is due to low coverage of traditional PAs, or the lack of sufficient data on biodiversity in the region.

### b. Appraisal of anthropogenic pressures

Reconciling biodiversity conservation concerns and the demands of economic development is among the greatest challenges of the 21^st^ century. We explicitly incorporated *Threats* (Fig. 2b) both, as factors that make conservation more costly, and also such that the final priority ranks represent such a reconciliation (Fig. 3). Across zones, a consistent pattern we observed was the compounding effects of agricultural expansion and urbanization, coupled with vegetation greening (indicative of increased year-round irrigation in agriculture areas) and linear infrastructure (Fig. 2b). Thus, the agricultural belts of (i) the northern semi-arid zone, (ii) the lowland plains of Northeast India, (iii) western and southern parts of the Deccan Peninsula, and urban hotspots representing major cities were ranked the highest in terms of anthropogenic pressures (Fig. 2b; Fig. S1). Of these, the northern semi-arid zone and the western parts of Deccan peninsula also had pronounced signatures of vegetation browning and greening, respectively. Vegetation browning could arise from forest degradation, deforestation, or forest fires; it could also be due to natural shifts in woody vegetation to grass dominated systems, or loss of green foliar biomass due to climate change impacts. Vegetation greening could be due to ecological restoration or natural regeneration of native species; but the more ubiquitous source is CO2 fertilization and the proliferation of invasive alien species (e.g., *Prosopis*, *Lantana*, etc.; Krishnaswamy et al. 2014; Parida et al. 2020; Piao et al. 2020).

**Figure 3.**
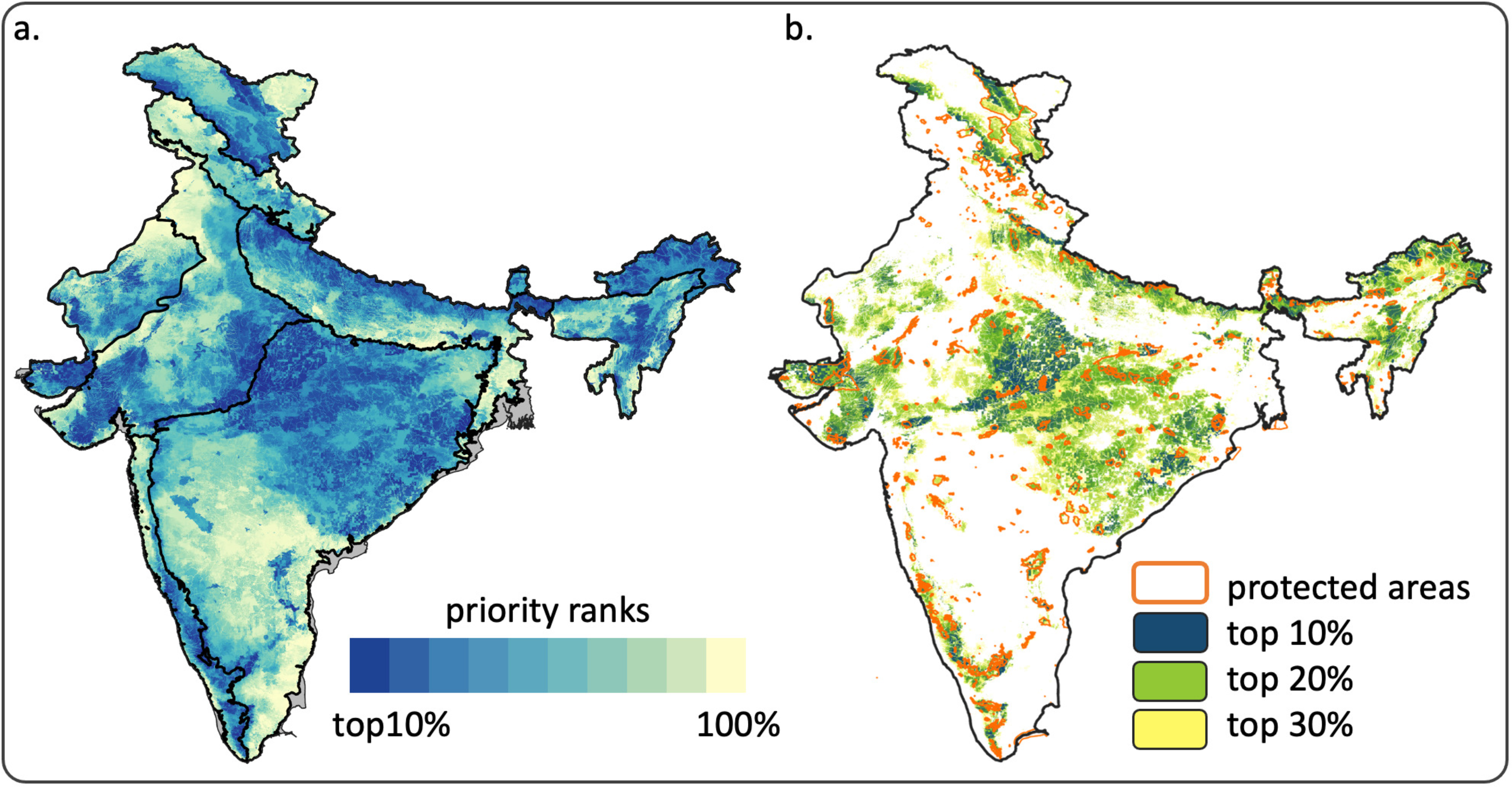
(a) Conservation priority ranks for the country, computed zone-wise and combined; (b) Top 30% priority areas from each zone demarcated as a set of three 10% blocks. Designated Protected Areas are overlaid to show the extent of overlap and spatial congruence with high priority locations.

Our results pertaining to threats, when viewed in the context of projected human population growth and the demands of meeting future food security concerns, are representative of areas that either have or will soon surpass thresholds beyond which interventions for sustainable land-use practices may not be feasible (Martin et al. 2022). On the other hand, our analyses could not fully elicit the broader deleterious impacts of certain human activities. These include threats that are (i) rapidly evolving in terms of scale, extent, and impacts, like hydropower dams and road networks (see Pandit & Grumbine 2012; Nayak et al. 2020), (ii) peculiar to certain regions of the country (e.g., expansion of oil palm plantations; Srinivasan et al. 2021), or (iii) difficult to quantify in terms of their long-term consequences (e.g., loss of connectivity; Vasudev et al. 2021). While we did incorporate projected temperature increase and future rainfall anomalies to account for climate change impacts, zones that are likely to be the most vulnerable to these threats, i.e., coastal areas and island systems, were not part of our assessment. Nevertheless, climate change is still projected to impact various ecosystems considered in our analysis in significant ways. For example, predicted increases in the magnitude and frequency of major floods can shape the distribution of biodiversity on the floodplains of Northeast India (see Goswami et al. 2021). Such influences underscore the importance of spatial conservation planning based on prioritization efforts of the kind we undertake here. We also note that climatic influences on terrestrial systems are extremely dynamic and are pivotally linked to national and global policy changes in the future.

### c. Landscape-level approach to conservation

Recent conservation literature has highlighted the importance of viewing shared habitats as coupled human–natural systems that are encompassed within ‘conservation landscapes’ (Das et al. 2006; Rodrigues et al. 2021; Vasudev et al. 2021). Across zones, we found that most designated PAs, which constitute ~5% of India’s land area and span an average size ~300 km^2^ (Ghosh-Harihar et al. 2019), were included as priority sites. But these were embedded within larger landscapes that also included high priority non-PA locations; contrary to our expectation that PAs would be over-represented in our results, 85% of the top 30% priority sites were outside PAs (Fig. 3). This was reinforced by the specific inclusion of ecosystems (such as the Open Natural Ecosystems referred to above) that are not part of India’s PA network, and sites critical for the supply of ecosystem services. Interestingly, locations that constituted the top 30% priority ranks in our study were largely connected (Fig. 3), even when we did not explicitly impose connectivity parameters via the prioritization model. To examine this further, we generated the ‘clumpiness’ index, which compares priority adjacencies with what would be expected at random (Hesselbarth et al. 2019). The index ranges from −1 (disaggregated) to +1 (clumped). When examined at the countrywide level, the index value was 0.80; within-zone analyses produced index values ranging from 0.77 to 0.88–– indicating high aggregation of priority sites at both spatial scales. Although our zone-wise analyses allowed for better geographic representation of sites, it still indicated lack of contiguity across some zone boundaries (see Fig. 3a). These aspects together emphasize that functional connectivity is an important consideration while implementing landscape-scale conservation interventions within our priority sites (see Brennan et al. 2022).

Traditional PAs, which typify a *land-sparing* approach to conservation, are mostly focused on forested habitats in the country. A substantially large proportion of biodiversity continues to inhabit unprotected, human-use landscapes, warranting a *land-sharing* approach. Realizing conservation and human well-being goals in the landscapes that encompass the 30% priority areas will therefore necessitate invoking Other Effective area-based Conservation Measures (OECMs). The core tenets of these approaches hinge fundamentally on effective and *equitable* models of governance that are fully cognizant of the complexities of socio–ecological systems, all of which are only beginning to be recognized in environmental and conservation policy (Alves-Pinto et al. 2021; Jonas et al. 2021). In India, this may be achieved through implementation of existing frameworks, for instance, in locations where communities are granted Community Forest Rights, and declaring areas as Critical Wildlife Habitats under the Forest Rights Act –– provisions which, at present, remain extremely under-utilized. Our spatially explicit landscape approach that considers linked biodiversity–ecosystem service dimensions can only succeed if the beneficiaries of ecological restoration, and those who may be otherwise impacted (typically the marginalized sections of society) are addressed in policies and implementation.

### d. Synergies and trade-offs within and among themes

In the final prioritization assessment, we assigned equal weightage to all input themes (equal but negative weight for the *Threats*). We ascertained the synergies and trade-offs between themes by examining alternative scenarios where (i) areas with high human impacts were prioritized rather than penalized, i.e., *Threats*-focused assessment; and (ii) *Habitats*, *Ecosystem Services* and *Biodiversity* were each iteratively afforded higher weightage compared to the others (see the Methods section for details). While there was reasonable concordance from our balance scenario (themes with equal weights) and those from individual theme-focused scenarios, some locations showed the stark mismatches indicative of the trade-offs (see Figures S3 to S5 in Supplementary File 1). These trade-offs may reflect regional peculiarities; biodiversity conservation could constrain provisioning services such as non-timber forest products or livestock-grazing in certain locations (like PAs). Increase in tree cover through habitat (mis)management could alter hydrologic services (see Joshi et al. 2018). Such trade-offs could also be spatially asynchronous––up-stream water abstraction in rivers, and disruption of sediment transport by dams can bear deleterious impacts on aquatic biodiversity and ecosystem services downstream. All these cases collectively highlight the problems with prioritizing areas based either on singular themes, or biasing prioritization on one theme over the other(s). Of course, there is also a trade-off in choosing locations that are of high conservation value and relatively secure (avoiding regions of high *Threats*) or those that are vulnerable (prioritising *Threats*). Ideally, the former is suited for interventions involving preservation (e.g., reserve design, OECMs), and the latter, interventions involving mitigation or restoration (e.g., mitigation of linear infrastructure impacts).

Besides the differences *between* themes described above, we acknowledge that there could be trade-offs even *within* the thematic dimensions considered here. For instance, we found very low spatial concordance between carbon versus blue water flux, and marginally higher correlation for carbon versus green water flux; the spatial mismatch was more pronounced at locations where values of above- and below-ground carbon reached very high levels (see Fig. S6 in Supplementary File 1). This relationship is not unique to our study area; carbon sequestration and hydrologic services do not necessarily work in synergy (Chisholm 2010). Regulatory services, like hydrologic or water services, can be generated only at certain size of catchment depending on factors such as climate, soils, geology and vegetation type. And while there could be synergy between carbon and water services to people at a local or regional scale (Clark et al. 2021), the transpiration from a large patch of forest can increase rainfall in other regions and benefit agriculture and communities elsewhere (Paul et al. 2018). These considerations of synergies and trade-offs are particularly relevant when viewed in the context of drastic biodiversity declines documented in the recent Living Planet Report (WWF 2022). The report reiterates the important links between biodiversity loss, climate change, ecosystem services (water) and food security–– aspects addressed in this study through our multi-criteria assessment and allocating equal importance (weights) to the constituent input attributes.

### e. Linking landscape prioritization and administration

Overlaying administrative (district) boundaries, we highlight 338 districts that play a key role in maintaining India’s biodiversity and ecosystem services (Fig. 4a and 4b); of these, 169 are ‘high priority’ districts, where natural habitats, biodiversity and ecosystem services are currently at optimal levels, and span a large spatial extent. The next 169 are ‘potential priority’ districts (Fig. 4, centre panel), where the three aspects are currently at relatively sub-optimal levels in terms of the extent of coverage. At this point, our aim was to also link our results with India’s aspirational districts, identified by the Indian government for economic development (i.e, the NITI Aayog aspirational districts; see Methods for details).

**Figure 4.**
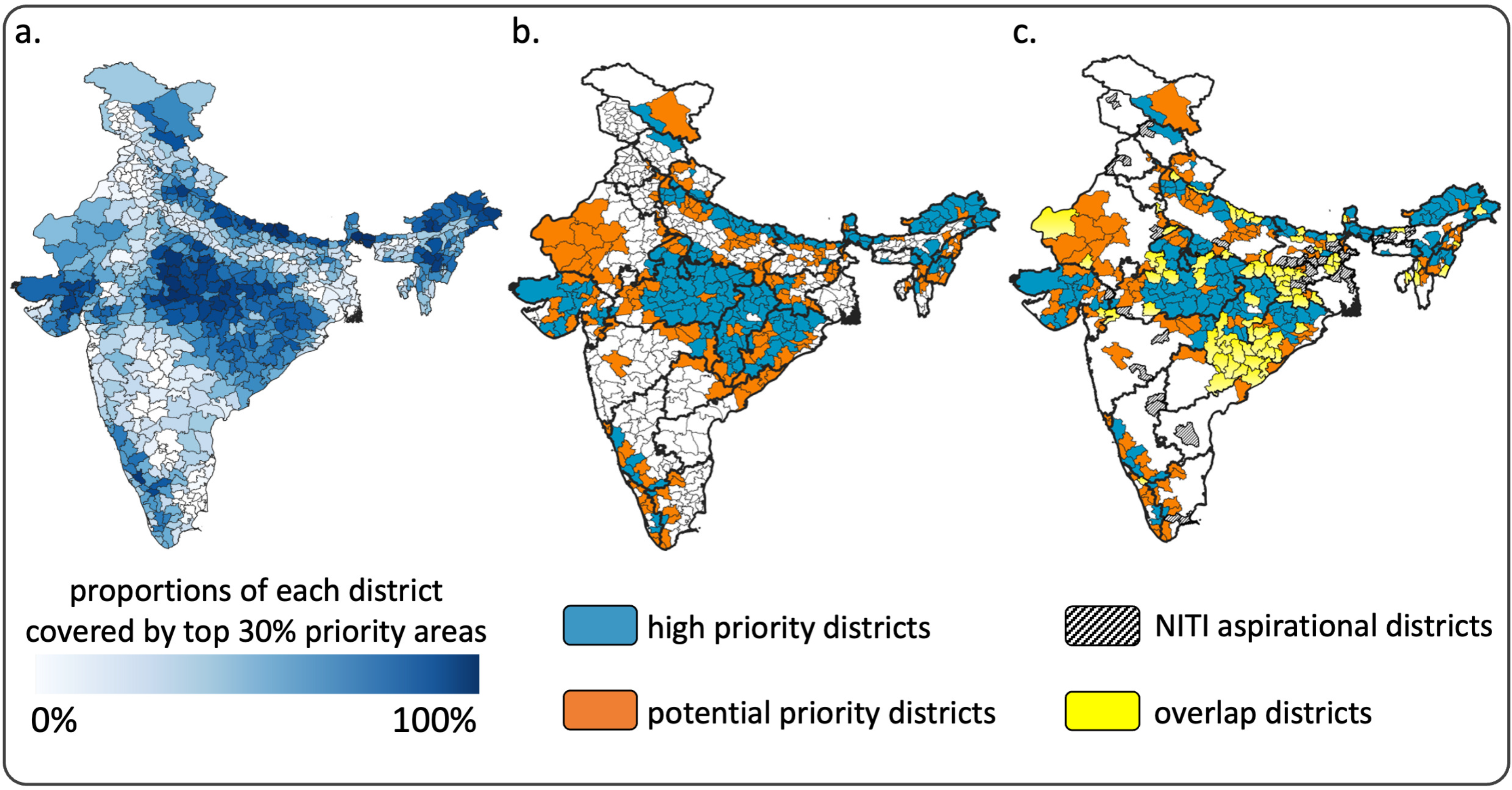
Results from spatial prioritization analyses processed at the scale of administrative units in the country. (a) Proportions of each district covered by areas identified as top 30% priority locations; (b) Districts in the top quantile (25%) identified as 169 ‘high priority’ districts and the next 25% quantile identified as 169 ‘potential priority’ districts; (c) NITI Aayog aspirational districts (112) earmarked by the Government of India, and their overlap with 338 priority districts identified in this study (72 districts overlap). Darker lines in (b) and (c) denote state boundaries.

Considered in conjunction with our results, we recommend that locations where the NITI aspirational districts overlap with ‘high priority’ districts (Fig. 4c), the management focus needs to be on retention of habitats, ecosystem services and biodiversity through both, state-driven and participatory approaches. This will require deprioritizing mega-infrastructure projects while promoting sustainable models of nature protection besides demarcation of PAs. Such approaches may entail community stewardship for biodiversity protection, co-management of habitats outside PAs, and nature-friendly livelihood development, within larger conservation landscapes. In locations where aspirational districts overlap with ‘potential priority’ districts, the management focus, besides retention of important sites, should also aim for proactive rewilding and ecological restoration efforts. Here, targeted actions are essential to mitigate negative impacts of infrastructure development through economically incentivized instruments like paying for ecosystem services, promoting agroforestry, and where appropriate, demarcating conservation and/or community reserves. These localized efforts and interventions need to be synergistically integrated within district- and state-level plans to ensure tangible impacts. For further deliberation, we provide maps showing spatial overlaps between NITI aspirational districts and (i) top 30% priority areas and (ii) threat ranks across the country (see Fig. S7 in Supplementary File 1).

### f. Practical considerations for on-ground implementation

To the best of our knowledge, this study represents the first countrywide assessment that attempts to link eco-socio-administrative dimensions for landscape-level prioritization in India. But we also submit that the spatial scale and resolution at which we make this assessment was constrained by several limitations associated with data availability that are commonplace across developing countries as well as global-level datasets alike. Our metric of species diversity, for instance, relied on range maps derived from IUCN, which do not have uniform accuracy across species (Ramesh et al. 2017; Srivathsa et al. 2020). In fact, range maps for many rare, endemic and data-deficient species in India were either unavailable, or, excluded from our assessment because they were completely inaccurate. We also used only a subset of ecosystem services deemed important for human well-being; services like pollination, forest produce extraction, rangeland services, or freshwater dependence, could not be included due to unavailability of spatially explicit data. Incorporating these features (when such information becomes available) may yield a more accurate and comprehensive assessment of priority locations and landscapes.

The relationship between ecosystem services and poverty alleviation (especially in the aspirational and poorer districts of India as defined by the NITI Aayog discussed above) is also driven by political economy of negotiations between stakeholders and those who manage or regulate ecosystem services (Vira et al. 2012). A detailed understanding of the associated vulnerabilities is important to enable ecosystem services to benefit the poor. If not internalized, an ecosystem services approach to conservation planning may fail because of resistance from those who are excluded or those who stand to lose most from such undertakings. The actions discussed above will also be critical in other countries of the Global South, where a large proportion of the people are directly dependent on forests and other natural ecosystems for their livelihoods. The nuances and location-based differences in prescriptive actions we discuss here reiterate the importance of (a) spatially explicit prioritization assessments incorporating regional expertise on biodiversity and threats, and (b) mainstreaming biodiversity and ecosystem service goals into district and state management plans for socio-economic development. In this context, we posit that India’s ambitious endeavours on climate change mitigation through reversal of land degradation (26 million ha), the Green India Mission (Ravindranath & Murthy 2010), as well as renewable energy projects initiated under the Intended Nationally Determined Contribution (NDC) will require a degree of reorientation.

## Conclusion

The recognition of links between ecosystem integrity and human health have led to broad and ambitious goals under global targets such as the Post-2020 Global Biodiversity Framework. However, repercussions of the COVID-19 pandemic have underscored the wide geographical imbalance in the capacities, resources, and vulnerabilities of different countries to achieve such goals. The Global South, in particular, aims to conserve biodiversity and meet climate change goals under scenarios of high dependency of people on natural resources (Fedele et al. 2021), coupled with aspirations of economic progress, and better standards of human life. To address these expectations, national-scale prioritization exercises such as ours, which combine conservation priorities, human well-being indices, economic development and infrastructure considerations can guide countries and governments towards meeting international targets set forth for the next decade (e.g., Strassburg et al. 2020); linkages between prioritization exercises and existing government schemes and administrative boundaries are crucial in this regard (see Srivathsa et al. 2020; Belote et al. 2021). In the context of elevated competition for scarce land and water, exacerbated due to future climate change, the importance of our framework lies in its ability to objectively and effectively address trade-offs at the intersection of sustainable development goals, conservation of biodiversity and ecosystem services.

## Methods

### a. Framework for prioritization

The main goal of our study was to prioritize representative landscapes for biodiversity conservation, and securing of ecosystem services and human well-being targets, while balancing these with the economic aspirations of 1.4 billion people. Our prioritization exercise is part of a larger program aimed at mainstreaming biodiversity conservation into the discourse on development and human well-being –– the Government of India’s National Mission for Biodiversity and Human Well-Being (NMBHWB; Bawa et al. 2021). Recognizing the importance of delineating, characterizing, and evaluating functional landscapes, the NMBHWB set up a working group in 2020, which was tasked with deliberating on and determining landscape-level approaches for biodiversity conservation, keeping nature’s benefit to human well-being as the central theme. Through consultations with 42 experts from across the country, the working group first outlined the need for, salient features of, and the prerequisites for a priority-setting exercise. Subsequently, the working group engaged with 18 field and domain experts (authors of the present study) to undertake the prioritization exercise in India.

### b. Prioritization approach

We used a two-step prioritization approach (Fig. 1) to ensure representation of geographies and eco-regions across India. We first chose biogeographic zones (‘zones’) as described by Rodgers and Panwar (1988) as an appropriate level of primary spatial classification. This level of classification splits the country into zones that have some similarity of biogeography, while across zones, they capture diversity of species, ecosystems, and human–nature relationships. The country has 10 zones: (i) trans-Himalayas, (ii) Himalayas, (iii) deserts, (iv) semi-arid, (v) Western Ghats, (vi) Deccan plateau, (vii) Gangetic plains, (viii) Northeast India, (ix) coasts and (x) islands. We excluded coasts and islands from our analyses as we deemed them to be unique and requiring a separate treatment of biodiversity, ecosystem services and threats. The eight selected zones are shown in Fig. 1 (additional descriptions are in Table S1). We then identified three broad focus themes, each of which encompassed a set of criteria for prioritization.

i. We considered ecosystems or natural habitats that require inclusion in priority landscapes. These layers, with zone-specific sets of priority ecosystems, represented our *Habitats* theme.
ii. Recognizing the increasing emphasis on human–nature links and joint well-being of biodiversity and people, we considered *Ecosystem Services* as the second theme.
iii. Species diversity, biodiversity hotspots and locations of species population sources collectively formed the *Biodiversity* theme.

Lastly, we included a *Threats* theme to recalibrate conservation priorities based on spatial variation in anthropogenic pressures and impacts on natural ecosystems and biodiverse areas that provision essential ecosystem services. The full list of input layers is referenced in Fig. 1 and detailed in Table S2.

### c. Spatial prioritization

There are several approaches to conduct spatial prioritization analyses, depending on the objectives of the study, type(s) of spatial data available, and the optimization functions of interest, along with corresponding software programs, of which MARXAN and Zonation are the most widely used (Watts et al. 2009, 2021; Moilanen et al. 2014; Sierra-Altamiranda et al. 2020; Silvestro et al. 2022). Zonation takes information on multiple input features to produce conservation ranks across the entire area of interest. Depending on the metrics used and the analytical specifications, it is possible to use either program to yield comparable results (see Delavenne et al. 2011). We found Zonation to be well-suited for our analysis, because it allowed us to seamlessly combine feature data (0/1), with quantitative data on ecosystem services across large regions at a relatively fine spatial resolution. The program is also robust to differences in scale (range of values) across different input layers. The program’s algorithm follows an iterative process to rank cells (in our case, 1-km^2^ pixels) within the extent of interest, by excluding cells one at a time and measuring the collective conservation value of retained cells. In other words, cells with the least value are removed first and those with the highest values are retained till the end. The final output thus generated is an optimized map whose pixels are ranked based on relative priority values. For additional details see Moilanen et al. (2014). For each zone, we implemented a set of six prioritization analyses (four theme-wise runs and two composite runs) as explained below.

#### Habitats

We used data from Roy et al. (2015) with 156 land-cover classes across the country. We first reduced the data into 28 habitat classes by grouping similar land cover classes, broadly following the forest type classification of Champion and Seth (1968). For instance, land cover types labelled “Grassland”, “Manmade grassland”, “*Lasiurus-Panicum* grassland”, “*Cenchrus-Dactyloctenium* grassland”, “*Aristidia-oropetium* grassland” and “*Sehima*-*Dichanthium* grassland” were reclassified collectively as “Grasslands”. Data from 25-m^2^ pixels of the original dataset were resampled at a 1-km^2^ resolution; the value assigned to each 1-km^2^ pixel for each of the 28 habitat classes reflected the proportion of 25-m^2^ pixels of the corresponding class. For each zone, we further sub-selected habitat classes to retain (a) the dominant vegetation classes (covering >1% of the total zone area), and (b) select rare, vulnerable habitats, determined based on our experience and knowledge of the landscape(s). For example, we included wet grasslands in the Northeast zone due to its ecological and livelihood importance, even though this habitat class covered only 0.25% of the zone. The final number of habitat types chosen for each zone thus varied between three (Desert zone) and 10 (Western Ghats zone; Table S2). All habitats were assigned equal weights for prioritization. We chose the ‘Core Area Zonation’ cell removal rule as it prioritizes pixels with the most important yet rare features; this way, pixels that contained high proportions of rare habitats were retained even if the other zone-wide dominant habitats within these pixels were low.

#### Ecosystem Services

We identified and used three layers that collectively reflected key ecosystem services, for which we were able to obtain or generate spatially explicit metrics across the country. These included (i) blue water flux, or the amount of water available as surface water or as ground water to meet utilitarian needs and for ecological flows in rivers, (ii) green water flux, or the amount of water lost via surface evaporation or through transpiration, which contributes to multiple ecosystem services, from flood-control in very wet regions to recycling back as rainfall in other regions, and (iii) carbon stock, a harmonized metric of aboveground and belowground biomass carbon density calculated using woody plant, grassland, and cropland biomass (Table S2). While these do not represent a comprehensive set of ecosystem services, we believe they capture two critical policy mandates for India: water security and climate change mitigation. We assigned equal weights to all three layers and chose the Additive Benefit Function ‘ABF’ cell removal rule in Zonation, to ensure that locations with relatively high values of all three features are prioritized higher than areas where individual features or subsets of the three features had high values.

#### Biodiversity

India supports an extremely high diversity of wildlife (within and outside designated Protected Areas); most of these species are found in higher densities here than elsewhere across their range. In considering attributes that best represent biodiversity, our goal was two-fold. First, we wanted to include areas that support high populations of species, and second, those areas that support and adequately represent the wide *diversity* of species outside Protected Areas (PAs). We therefore used PAs, reasonably assuming that these locations currently harbor source populations of many species. In addition, we also used Key Biodiversity Areas (KBAs; BirdLife International 2021) to include locations outside designated PAs with high diversity of species that are of ecological or conservation importance (i.e., PAs and KBAs did not spatially overlap in our analysis and were 0/1 data). Next, we collated distribution ranges of imperilled mammals, birds, amphibians, and reptiles––species categorized as Critically Endangered, Endangered, Vulnerable or Threatened as per the IUCN Red List (IUCN 2020). We limited our criteria to only include species for which the range distribution data were reliable; for a subset of species, we also corrected and modified the IUCN maps (based on expert consultation) prior to inclusion in our analyses. For each zone, we stacked the species maps and generated a beta-diversity index, calculated as the Bray-Curtis distance between a focal cell and a hypothetical reference cell which had all threatened species of the corresponding zone. This gave us an index of zone-wise species turnover. We assigned equal weights to all three layers (treating PAs and KBAs with high population densities to be as important as the layer characterizing species turnover) and chose the ABF cell removal rule in Zonation to prioritize areas based on this theme.

#### Threats

We identified 11 attributes that negatively impact biodiversity, ecosystem services and habitats to varying degrees, and are likely to reduce the ease of implementing conservation actions. These included human population density, livestock population density, urbanization, linear infrastructure, mines, river fragmentation, agricultural expansion, vegetation greening and vegetation browning (both of which can have positive or negative impacts on ecosystem services and biodiversity), future climate warming and future rainfall anomaly (see Table S2 for details). We assigned zone-specific weights to these layers following a consensus-approach, based on the collective field knowledge of the assessors. The weights assigned to the threats varied across zones, reflecting the relative severity of impacts in the corresponding zone (see Table S3 for zone-wise threat ranks). We chose the ABF cell removal rule again, so that locations under more severe threat, and those under multiple threats were ranked high. This ranking thus combines information on the presence and intensity of threats across space.

#### Composite

In addition to the four sets of theme-wise analyses described above, we also undertook two composite analyses. First, we used the outputs from the theme-wise analyses of *Habitats*, *Ecosystem Services* and *Biodiversity* features. All input layers were assigned equal weights and the ABF cell removal rule was specified. Second, we used the aforementioned three layers along with the rank output from the *Threats* analysis. In this case, we assigned equal positive weights to *Habitats*, *Ecosystem Services* and *Biodiversity*, and an equal but negative weight to *Threats* (i.e., 1, 1, 1, and −1; see Wendland et al. 2010 and Koschke et al. 2012 for justification on using equal weights for input attributes). The first analysis thus gave us an assessment of conservation importance, while the second incorporated feasibility. To assess the sensitivity of our weighting scheme, we performed similar composite analyses to consider alternate scenarios where (i) the *Threats* layer was assigned a weight of +1, reflecting results such that areas with high human impacts are also prioritized rather than penalized, and (ii) the *Threats* layer was weighted −1, but the *Habitats*, *Ecosystem Services* and *Biodiversity* were each iteratively assigned higher weights than the other two, i.e., 1, 0.5 and 0.5. These outputs are presented in Supplementary File 1.

### d. Alignment with global and national policy

We collated results across zones post-prioritization to demarcate the top 30% priority areas in the country, so as to align our results with the Post-2020 Global Biodiversity Framework targets. The areas were selected based on pixel ranks in each zone; these covered 30% of the area in every zone, and therefore, collectively included 30% of the country (sans the coasts and islands). This approach ensured representation of unique features from each zone, encompassing a more diverse set of habitats, ecosystem services, biodiversity, and associated threats. We then overlaid administrative units (districts) and calculated the proportion of each district covered by pixels with the top 30% ranks. We chose two sets of districts: (1) those in the top 25% quartile, which we term ‘high priority’ districts, and (2) those in the subsequent 25% quartile, which we term ‘potential priority’ districts. We juxtapose these results with districts earmarked by the Government of India’s flagship Aspirational Districts Programme for economic development. India’s Aspirational Districts Program was launched in 2018 by the Government of India through the think tank, the National Institute for Transforming India (NITI) Aayog. The NITI Aayog replaced the erstwhile Planning Commission, with the aim of achieving sustainable development goals for the country. The program has earmarked over 100 of India’s most economically backward districts as ‘aspirational’ districts to help reduce regional imbalances in development. The program aims to reduce disparity across regions, and improve baseline rankings of district-level development using real-time data on 49 indicators across five thematic sectors including Health & Nutrition, Education, Infrastructure, Financial Inclusion, and Skill Development. Of particular relevance to our assessment is the program’s thrust towards increasing road connectivity and intensification of agriculture (Sarkar et al. 2022; additional details available at www.niti.gov.in/aspirational-districts-programme). Our maps were generated with the goal of directly offering prescriptive management actions for the governments of the corresponding administrative units.

## Data availability

All the analyses are based on open-source datasets; details are provided in Supplementary Table S2.

To whom correspondence should be address: Arjun Srivathsa

## Acknowledgements

This work was catalysed and supported by the Office of the Principal Scientific Adviser to the Government of India as part of the National Mission for Biodiversity and Human Well-Being. A.S. was supported by the Department of Science and Technology–Government of India’s Innovation in Science Pursuit for Inspired Research (DST INSPIRE) Faculty Award. We are grateful to J. Zacharias, V. Athreya, P. Bindra, A. Harihar, U. Ganguly, A. Kumar, M. Manuel, S. Babu, A. Kshettry, S. Nijhawan, M. Muralidharan, N. Namboothiri, Y. Jhala, K.S. Subin, S. Datta, M. Sen, S. Madhulkar, T. Thekaekara, V. Vasudevan, I. Anwardeen, S. Sahu, S. Reddy, V. Varadhan, K.T. Subramaniam, C. Madegowda, C. Meenakshi, K. Karanth, P. G. Krishnan, T. Dash, A. Chanchani and A. Bijoor for partaking in discussions on landscape-scale conservation in India. We thank K. Bawa and R. Chellam for their guidance, R.G. Rodrigues, A. Samrat and N. Srinivas for assistance with data processing and compilation, and the National Centre for Biological Sciences-TIFR and Ashoka Trust for Research in Ecology and the Environment (part of the Biodiversity Collaborative) for facilitating this study.

## Author contributions

All the authors were involved in the conceptualization of the study. AS, DV, TN led the analysis; AS, DV, TN, AD, RN and SV processed and analyzed the data. AS, DV, VRG, AN, JK and UR prepared the first draft. All authors provided critical feedback and approved the final version of the manuscript.

