## Supplementary File 1 for "Prioritizing landscapes to reconcile biodiversity conservation, ecosystem services, and human well-being in India"

**Figure S1. (a) spatial overlap of the top 30% pixel ranks showing (i) ‘HB + BD’– combined habitats and biodiversity (ii) ‘only ES’– only ecosystem services (iii) ‘HB–BD–ES’– habitats, biodiversity and ecosystem services; (b) regions and major cities with the highest ranks for Threats in each biogeographic zone.**

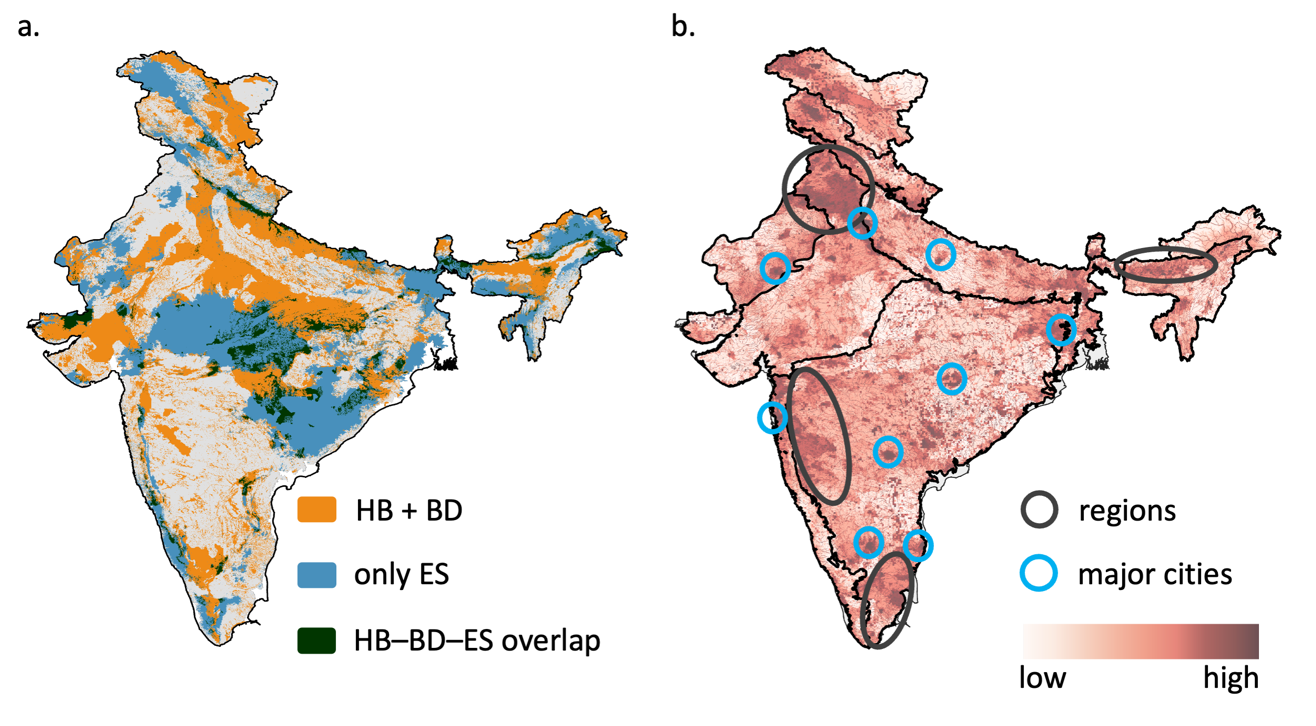

**Figure S2. Alternative scenario where the *Threats* theme was assigned a weight of +1, reflecting results such that areas with high human impacts are prioritized rather than penalized (Habitats=1, Ecosystem Services=1, Biodiversity=1, and Threats=1). (a) Pixel-wise difference in between priority ranks of this alternative scenario and priority ranks from the ‘Balanced scenario’ (Habitats=1, Ecosystem Services=1, Biodiversity=1, and Threats= -1). White tones indicate locations with synergy, colored tones (green/purple) indicate locations with trade-offs; green indicates locations ranked high under the *Threats* scenario, but not under the *Balanced* scenario, while purple indicates the opposite; (b) Scatterplot of 1,000 randomly selected pixel-wise values between *Threats*-focused prioritization and Balanced prioritization.**

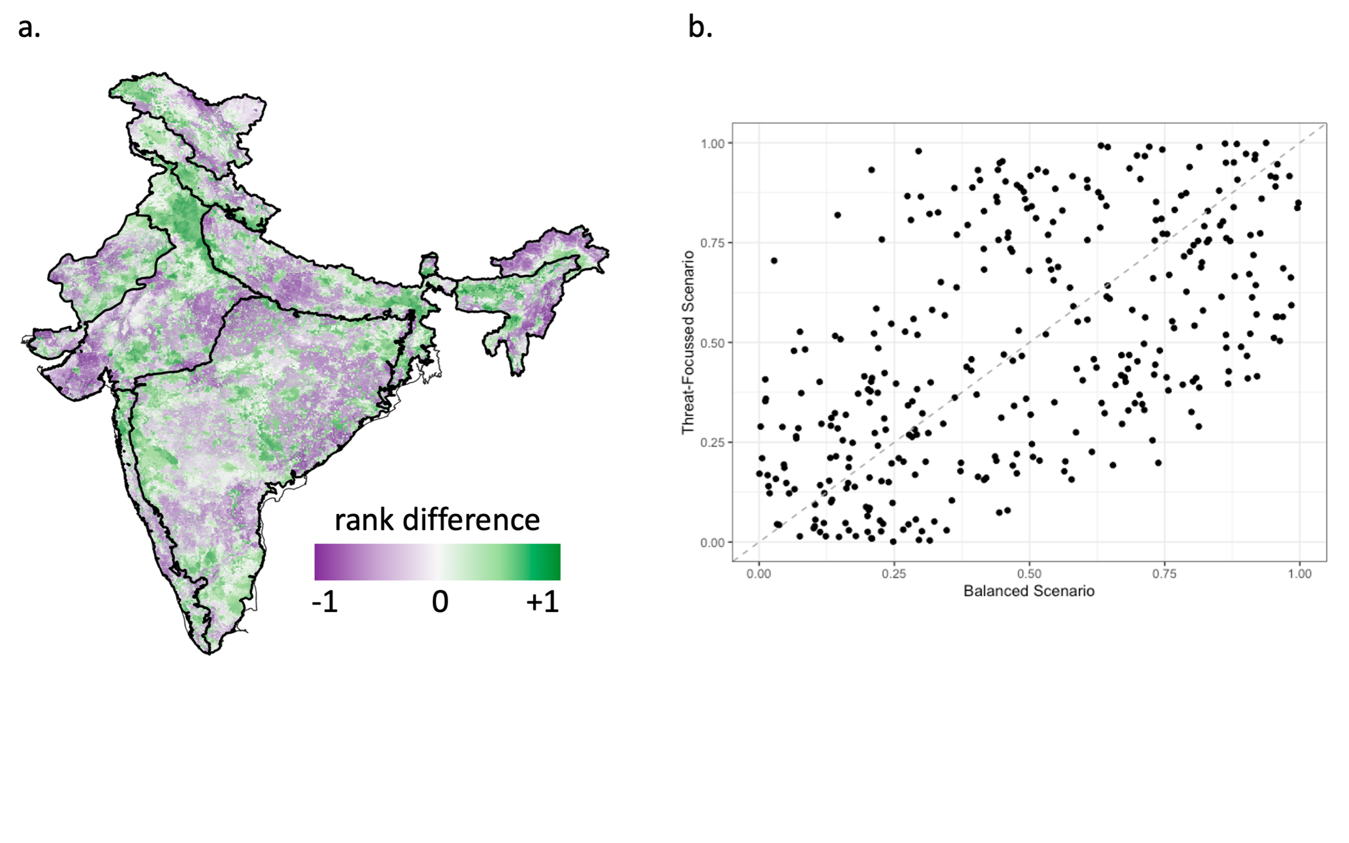

**Figure S3. Alternative scenario with *Habitats* theme-focused prioritization (Habitats=1, Ecosystem Services=0.5, Biodiversity=0.5, and Threats= -1). (a) Pixel-wise difference in between priority ranks of the *Habitats*-focused alternative scenario and priority ranks from ‘Balanced scenario’ (Habitats=1, Ecosystem Services=1, Biodiversity=1, and Threats= -1). White tones indicate locations with synergy, colored tones (green/purple) indicate locations with trade-offs; green indicates locations ranked high under the *Habitats* scenario, but not under the *Balanced* scenario, while purple indicates the opposite; (b) Scatterplot of 1,000 randomly selected pixel-wise values between *Habitats*-focused prioritization and Balanced prioritization.**

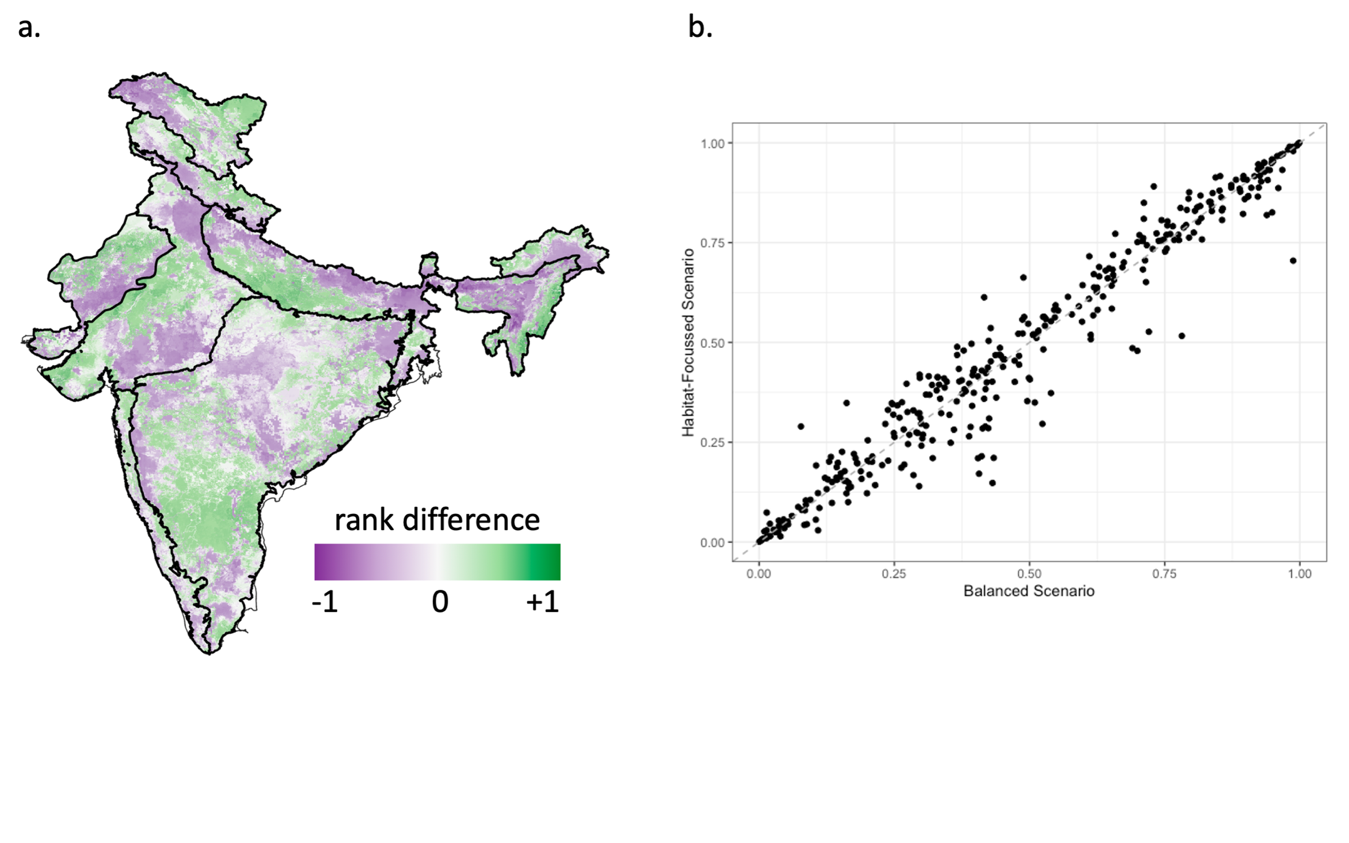

**Figure S4. Alternative scenario with *Ecosystem Services* theme-focused prioritization (Habitats=0.5, Ecosystem Services=1, Biodiversity=0.5, and Threats= -1). (a) Pixel-wise difference in between priority ranks of the *Ecosystem Services* alternative scenario and priority ranks from ‘Balanced scenario’ (Habitats=1, Ecosystem Services=1, Biodiversity=1, and Threats= -1). White tones indicate locations with synergy, colored tones (green/purple) indicate locations with trade-offs; green indicates locations ranked high under the *Ecosysted Services* scenario, but not under the *Balanced* scenario, while purple indicates the opposite; (b) Scatterplot of 1,000 randomly selected pixel-wise values between *Ecosystem Services*-focused prioritization and Balanced prioritization.**

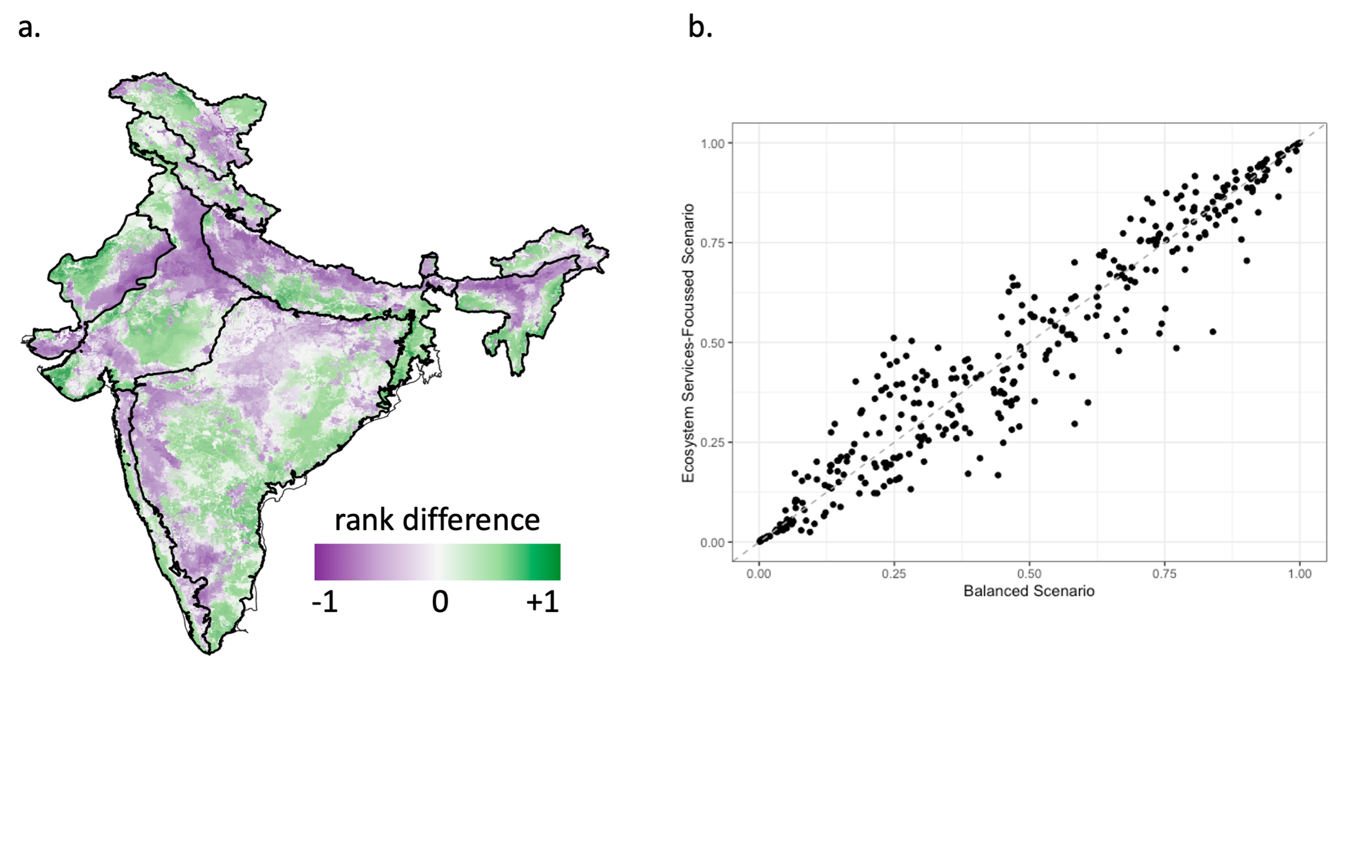

**Figure S5. Alternative scenario with *Biodiversity* theme-focused prioritization (Habitats=0.5, Ecosystem Services=0.5, Biodiversity=1, and Threats= -1). (a) Pixel-wise difference in between priority ranks of the *Biodiversity* alternative scenario and priority ranks from ‘Balanced scenario’ (Habitats=1, Ecosystem Services=1, Biodiversity=1, and Threats= -1). White tones indicate locations with synergy, colored tones (green/purple) indicate locations with trade-offs; green indicates locations ranked high under the *Biodiversity* scenario, but not under the *Balanced* scenario, while purple indicates the opposite; (b) Scatterplot of 1,000 randomly selected pixel-wise values between *Biodiversity*-focused prioritization and Balanced prioritization.**

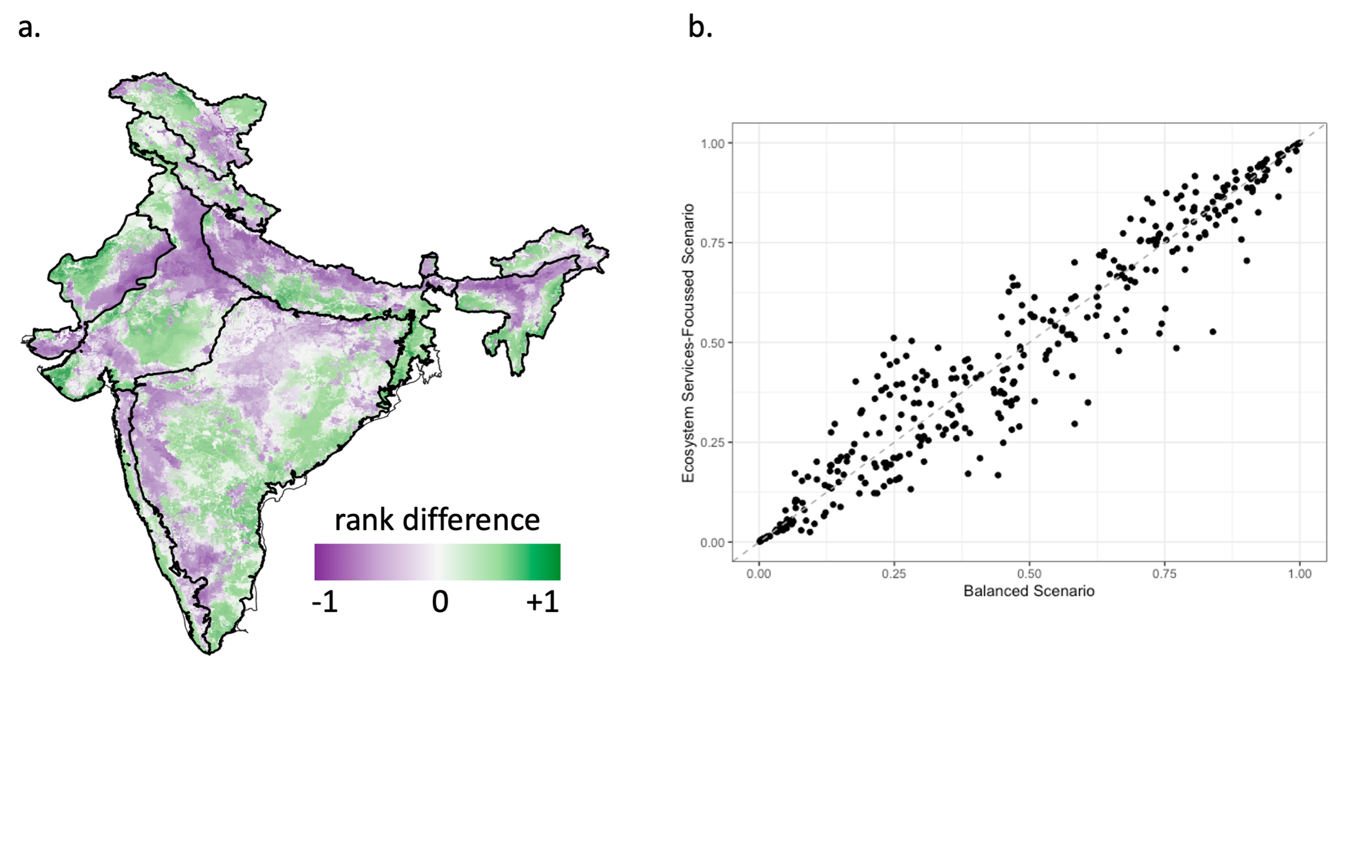

**Figure S6. Scatterplots showing the spatial concordance between carbon stock and (a) blue water flux, and (b) green water flux. Dots represent individual pixel-level values from across the country, with a sub-sample of 1000 points selected randomly and re-scaled for ease of depiction. Spearman rank correlation between carbon and blue water flux = 0.2; Spearman rank correlation between carbon and green water flux = 0.5.**

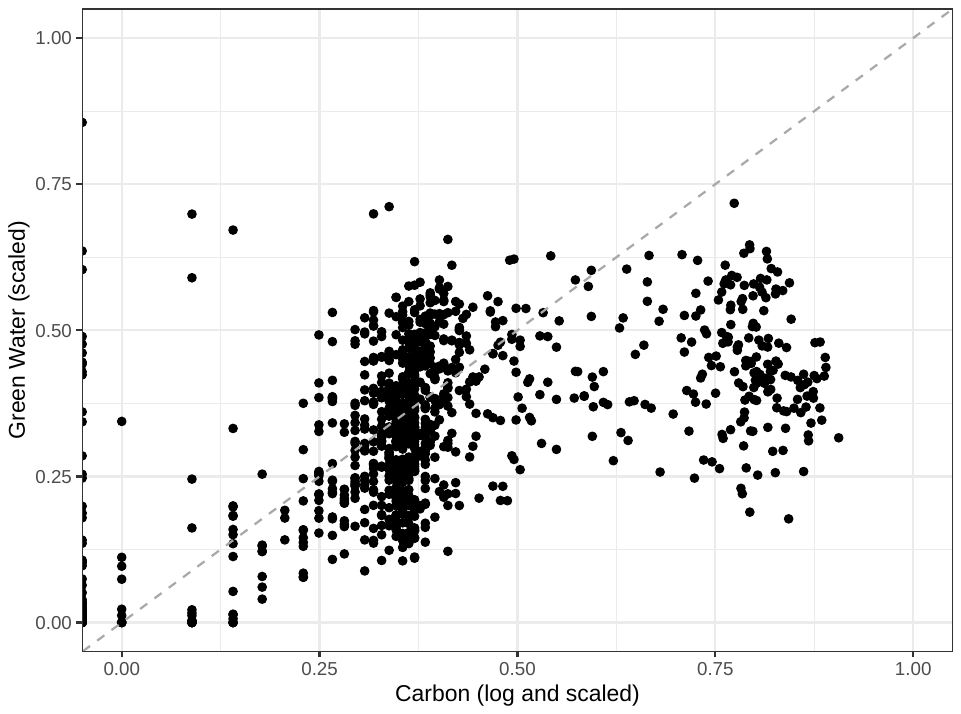
 **a.	b.**

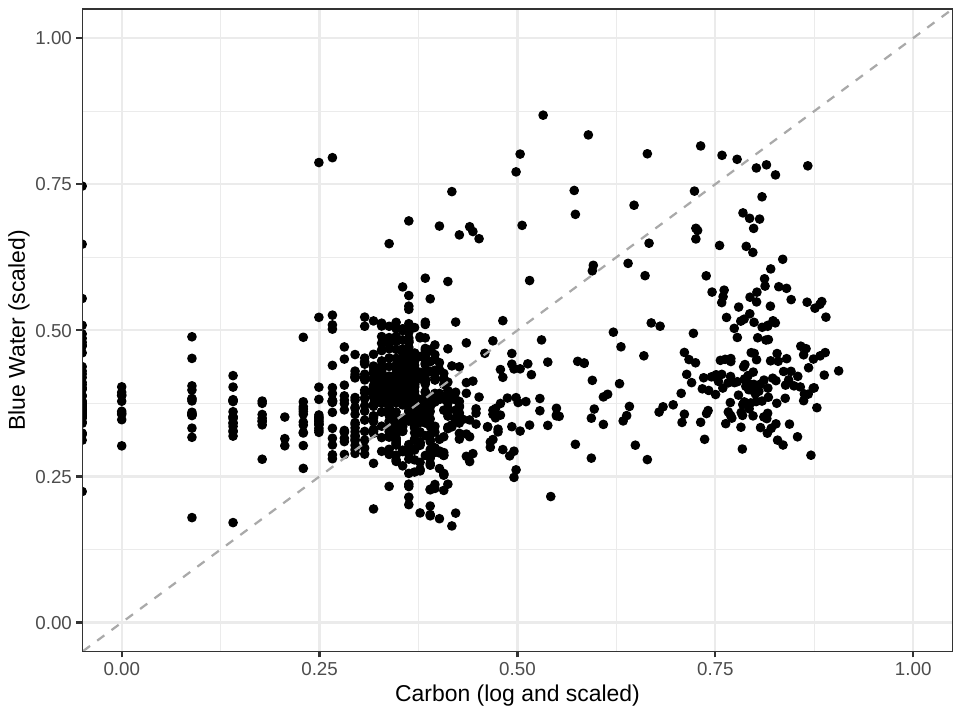

**Figure S7. (a) Top 30% priority areas from each zone demarcated as a set of three 10% blocks. NITI Aayog districts are overlaid on the top 30% priority areas to depict the extent of overlap; (b) NITI Aayog districts overlaid on map with threat ranks across all biogeographic zones.**

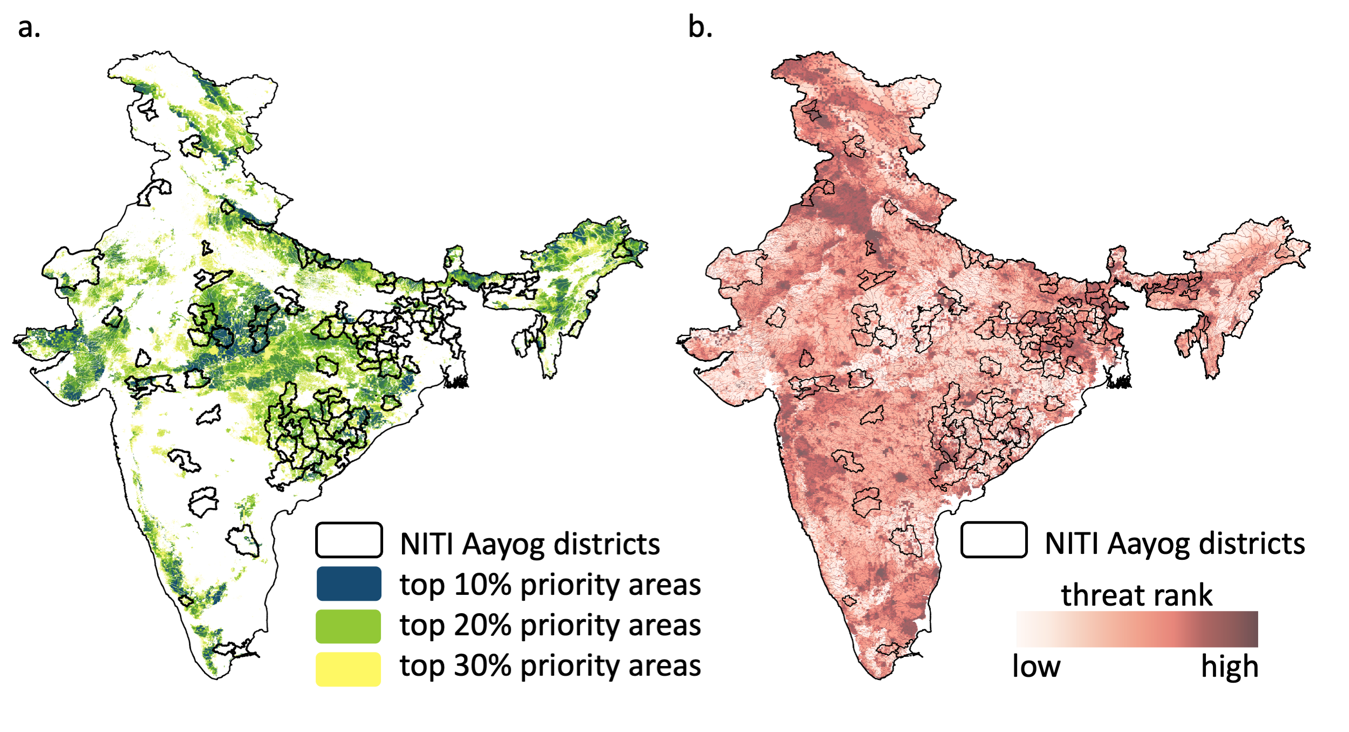
