## Supplementary material for "Prioritizing landscapes to reconcile biodiversity conservation, ecosystem services, and human well-being in India": Table S1

**Table S1.** Eco-climatic characteristics of the 8 biome-zones considered in this study. Classification of the biomes are as described by Rodgers and Panwar (1988) This level of classification splits the country into zones that have some similarity of biogeography, while across zones, they capture diversity of species, ecosystems, and human–nature relationships.

| **Sl.** | **Biogeographic Zone** | **Provinces** | **Total Area (km2)** | **Dominant Habitats** | **Proportion area under PAs** | **Human Population**  **(in mln)** | **Description** |
| --- | --- | --- | --- | --- | --- | --- | --- |
| 1. | Trans Himalayas | a. Ladakh Mountains  b. Tibetan Plateau | 182, 293 | a. Moist-Alpine Scrub b. Dry-Alpine Scrub c. High Altitude plains d. Open Habitats | 24.1% | 1.55 | **Climatic characteristics:** cold and arid  **Physical Attributes:** Altitude range 4500 – 6000 metres. Extensive areas consists of bare rock and glaciers.  **Ecological importance:** Large assemblage of mammals like wild ungulates, snow leopard, black and brown bears, ibex, grey wolf and the migratory black necked cranes. |
| 2. | Himalayas | \| a. North-West Himalaya b. West Himalaya c. Central Himalaya d. East Himalaya | 218, 874 | a. Tropical Wet Evergreen b. Subtropical Broad leaved c. Montane Wet Temperate d. Himalayan Moist Temperate e. Tropical Semi Evergreen f. Tropical Moist Deciduous g. Tropical Dry Deciduous and tropical dry forests h. Tropical Thorn i. Subtropical Pine j. Moist-Alpine scrub k. Dry-alpine scrub l. High altitude plains m. Open habitats | 11.4% | 29.4 | **Climatic characteristics:** A large gradient of temperature and moisture from alpine to tropical climatic regions  **Physical Attributes:** High altitude and steep gradient.  **Ecological importance:** Tropical rainforests in the East and subtropical and alpine forests in the Central and Western Himalayas. Key large mammals include wild sheep, ibex, musk deer, red pandas, dholes and leopards. |
| 3. | Deserts | a. Thar  b. Katchchh | 214, 091 | a. Tropical thorn b. Grassland c. Open habitats | 5.4% | 23.58 | **Climatic characteristics:** Arid bioclimate with very hot dry summers and very cold winters. Annual rainfall about 200 mm largely restricted to July-October, with frequent droughts.  **Physical Attributes:** Thar is the only dry desert in India, comprising plains interspersed with small sand dunes. Kutch is a large broken plain with dry open habitats, bounded by the coast to the south and Rann of Kutch to the north- a large seasonally inundated salt marsh.  **Ecological importance:** Plants are mostly xerophytic. Animals found in this region include the Great Indian bustard, lesser florican, migratory Macqueen's bustard, sandgrouses, coursers, raptors, chinkara, desert fox, and spiny-tailed lizard among others. |
| 4. | Semi-arid | a. Punjab Plains  b. Gujrat Rajputana | 532, 715 | a. Tropical Moist Deciduous b. Tropical Dry Deciduous c. Tropical thorn d. Ravines* e. Tropical Dry Forests f. Open Habitats | 3.5% | 221.47 | **Climatic characteristics:** Semi-arid bioclimate with a long and pronounced dry period, comprising hot and dry summers, cold winters, and annual rainfall of about 400 mm in short spell (July - October).  **Physical Attributes:** It is a transitional zone between the deserts in the West and the forests of Central India. Includes Aravalli and Vindhyan mountain ranges. It has a discontinuous vegetation cover with open habitats and soil-water deficit throughout the year.  **Ecological importance:** The vegetation includes discontinuous thorn forests, savannah, grasslands and scrub, and is highly fragmented by agriculture. It is home to species like the Great Indian bustard, lesser florican, quails, sandgrouses, coursers, chinkara, Indian fox, wolf and hyaena. |
| 5. | Western Ghats | a. Malabar Plains  b. Western Ghats Mountains | 131, 430 | a. Tropical Wet Evergreen b. Subtropical broad leaved Hill c. Tropical Semi Evergreen d. Tropical Moist Deciduous e. Tropical Thorn f. Littoral and Swamp g. Tropical Dry Forests h. Grassland* i. Riverine | 11.1% | 52.87 | **Climatic characteristics:** The mountains intercept monsoon winds from the southwest and create a rain shadow in the region to their East.  **Physical Attributes:** Average altitude of 900–1500 metres above sea level.  **Ecological importance:** It is a globally recognized biodiversity hotspot and has high levels of endemism. |
| 6. | Deccan plateau | a. Central Highlands  b. Chhota Nagpur  c. Eastern Highlands  d. Central Plateau  e. Deccan South | 1, 379, 961 | a. Tropical Semi Evergreen b. Tropical Moist Deciduous c. Tropical Dry Deciduous d. Tropical Thorn e. Tropical Dry Evergreen* f. Tropical Dry Forests g. Grassland* h. Open Habitats i. Riverine | 3.7% | 414.7 | **Climatic characteristics: Varies widely from** semi-arid parts in the rain shadow of the Western Ghats, to the wooded savannahs of central Indian highlands with brief periods of rain and a prolonged cold-dry and hot-dry periods.  **Physical Attributes:** It is the largest unit of the Peninsular Plateau of India. It has an average altitude of 600 metres and comprise of different types of forests.  **Ecological importance:** Grazing animals like the blackbuck and chinkara are seen in abundance here, among vegetation that mainly comprises short trees and shrubs. Globally recognized important landscapes for tiger populations in India. |
| 7. | Gangetic Plains | a. Upper Gangetic  b. Lower Gangetic | 355, 042 | a. Tropical semi evergreen* b. Tropical moist deciduous c. Tropical dry deciduous, tropical thorn, and tropical dry forests d. Littoral and Swamp e. Grassland* f. Wet grasslands* g. Open habitats h. Riverine i. Wetlands | 2.4% | 376.27 | **Climatic characteristics:** Varies from arid regions in the west to hot-humid zones in the eastern parts.  **Physical Attributes:** The Gangetic Plain in North India extends up to the foothills of the Himalayas. Most parts are flat, and most of the region includes agriculture land.  **Ecological importance:** High human population that is dependent on agro-based economy. Dominant trees in the forests are teak (*Tectona grandis*), sal (*Shorea robusta*), rosewood, mahua (*Madhuka indica*)and khair (*Acacia catechu*). |
| 8. | Northeast india | a. Brahmaputra Valley  b. North-East Hills | 171, 382 | a. Tropical Wet Evergreen b. Subtropical Broad Leaved Hill c. Tropical Semi Evergreen d. Tropical Moist Deciduous e. Tropical Thorn f. Subtropical Pine g. Tropical Dry Forests h. Grassland i. Wet Grassland* j. Open habitats k. Riverine l. Wetlands* | 4.4% | 46.38 | **Climatic characteristics:** Northeast India has a subtropical climate with high rainfall.  **Physical Attributes:** Largely hilly and mountainous region with altitudes ranging from 50–7000 metres.  **Ecological importance:** Its forests support several species of orchids, bamboos, ferns, and mammals like hoolock gibbon and tiger. |

*These habitats are not dominant, but were identified as rare or vulnerable in each biome.
