## Supplementary material for "Prioritizing landscapes to reconcile biodiversity conservation, ecosystem services, and human well-being in India": Table S2

**Table S2.** Details and description of data used as input attributes for zone-wise prioritization analyses in India. The table also includes information on the data source, spatial resolution and the corresponding year from which the data were sourced.

| **Theme** | **Layer name** | **Data description** | **Resolution at source** | **Data source** | **Year** |
| --- | --- | --- | --- | --- | --- |
| *Habitats* | **Land cover** | 153 land cover categories (from Roy et al. 2015) were combined/reduced to 28 classes, broadly following the forest type classification described by Champion and Seth (1968). | 25m | Indian Institute Remote Sensing, Govt. of India; http://bis.iirs.gov.in | 2015 |
| *Ecosystem services* | **Blue water flux** | Annual Blue Water Flux; Quantum of water that is available as surface and ground water | 1km | Rainfall: CHIRPS (Funk et. al. 2015) ET: USGS SSEBop Dataset (Senay et al. 2013: https://doi.org/10.1111/jawr.12057) | 2019 |
|  | **Green water flux** | Annual Green Water Flux; Quantum of water that is lost as evaporation from surface and as transpiration from plants | 1km | Rainfall: CHIRPS (Funk et. al. 2015) ET: USGS SSEBop Dataset (Senay et al. 2013: https://doi.org/10.1111/jawr.12057) | 2019 |
|  | **Carbon** | Harmonized metric of aboveground and belowground biomass carbon density calculated using woody plant, grassland, and cropland biomass | 300m | Spawn, S.A., and H.K. Gibbs. 2020. Global Aboveground and Belowground Biomass Carbon Density Maps for the Year 2010. ORNL DAAC, Oak Ridge, Tennessee, USA. https://doi.org/10.3334/ORNLDAAC/1763 | 2010 |
| *Biodiversity* | **Protected Areas** | Protected Areas (National Parks and Wildlife Sanctuaries) of India were downloaded in vector format from WDPA and updated using layers available from the Wildlife Institute of India. | Vectorfile | World Database on Protected Areas (WDPA), ENVIS Centre on Wildlife & Protected Areas (Wildlife Institute of India, WII); https://www.protectedplanet.net and http://wiienvis.nic.in/Home.aspx | 2019 |
|  | **Key Biodiversity Areas** | Key Biodiversity Areas; a request for database of south Asia region was made to and obtained from BirdLife International. The data include locations currently identified as "key biodiversity areas". | Vectorfile | World Database of Key Biodiversity Areas. March 2021 version. (BirdLife International 2021) | 2021 |
|  | **Threatened species** | Zone-wise beta-diversity of species; distribution maps of threatened species of mammals, birds, reptiles, and amphibians were compiled from IUCN Red List database and the beta-diversity calculated for each biogeographic zone at 7 sq.km spatial resolution. The beta-diversity index was calculated as the Bray-Curtis distance between a cell and hypothetical reference site which had all threatened species of the corresponding zone. | Vectorfile | The IUCN RedList of Threatened Species; https://www.iucnredlist.org/ | 2018 |
| *Threats* | **Human population** | Estimated human population for the year 2020 (extrapolated using baseline population from 2011 census by Govt. of India). Metric calculated as density of people per 1sq. km | 1km | NASA Socioeconomic Data and Applications Center (SEDAC); https://sedac.ciesin.columbia.edu | 2021 |
|  | **Livestock population** | Livestock (cattle, goats, sheep) population data compiled from government census records at the taluk (administrative unit) level. The metric reflects the density, calculated as number of livestock heads per taluk. | taluk (sub-district) | All India Livestock Census 2012, Govt. of India; http://dahd.nic.in | 2012 |
|  | **Urbanization** | Multi-temporal classification of built-up presence. Data coded as '0'- areas built-up before 1990, and '1'- areas built-up after 1990 (up to 2014). The metric is a surrogate for extent of urbanization | 30m | Global Human Settlement Layer (GHS-BUILT R2018A); https://ghsl.jrc.ec.europa.eu/ghs_bu2019.php | 1975–2014 |
|  | **Linear infrastructure** | Density of linear infrastructure (roadways and railways), downloaded as vectors and processed as length per 1 sq.km pixels across the country | Vectorfile | OpenStreetMap (OSM) Project; http://download.geofabrik.de | 2020 |
|  | **Mines** | Locations of mines and quarries across India were digitized. The vector file was converted into a raster of 1km resolution. All mines <1km size were upscaled to reflect one pixel. | Vectorfile | India Under Construction; https://indiaunderconstruction.com/: Nayak et al., 2020 (https://doi.org/10.1016/j.landusepol.2020.104619) | 2017 |
|  | **River fragmentation** | Database on global free flowing rivers was used to obtain information on connectivity status of rivers. The degree of connectivity was assessed at every 2 km stretch of each river. The index ranged from ‘0’- no connectivity to ‘100’- full connectivity | Vectorfile | Grill et al. 2019 | 2019 |
|  | **Agriculture expansion** | Time series of consistent global land cover maps collated on annual basis from 1992 to 2018. Data coded as '0'- land used for agriculture in 1992, and '1'- non agricultural land converted to agriculture between 1992 and 2018. The metric is a surrogate for agricultural expansion | 300m | European Space Agency-Climate Change Initiative: Land Cover; http://www.esa-landcover-cci.org | 1992–2018 |
|  | **Vegetation greening** | The metric represents agricultural intensification in areas under cultivation, and expansion of invasive plants in natural habitats. Senslope applied to time-series of annual ecoclimatic distance; NDVI3g | 9 km | Krishnaswamy et al. (2009) | 2009 |
|  | **Vegetation browning** | The metric represents reduction or drop in primary productivity. Senslope applied to time-series of annual ecoclimatic distance; NDVI3g | 9 km | Krishnaswamy et al. (2009) | 2009 |
|  | **Future climate warming** | Ensemble of Downscaled Outputs from 19 Global Climate Models (RCP 4.5) | 30 sec | WorldClim | 2045 |
|  | **Future rainfall anomaly** | Ensemble of Downscaled Outputs from 19 Global Climate Models (RCP 4.5) | 30 sec | WorldClim | 2045 |
