## Supplementary material for "Prioritizing landscapes to reconcile biodiversity conservation, ecosystem services, and human well-being in India": Table S3

**Table S3.** Ranks assigned to individual *Threat* attributes to reflect their relative severity in terms of impact on habitats, ecosystems services and biodiversity in each biogeographic zone. Details on the process involved in assigning the ranks are included below the table.

| **Zone** | **Threat Rank** | | | | |
| --- | --- | --- | --- | --- | --- |
|  | **1** | **2** | **3** | **4** | **5** |
| *Trans Himalayas* | Agricultural Expansion | Urbanization | Livestock Density | Future Climate Warming |  |
|  |  | Greening | Future rainfall anomaly |  |  |
|  |  | Browning | Mines |  |  |
|  |  | Human Density | River fragmentation |  |  |
|  |  |  | Linear Infrastructure |  |  |
| *Himalayas* | Agricultural Expansion | Livestock Density | Human Density | Future rainfall anomaly | Future Climate Warming |
|  |  | Urbanization | Greening | Linear Infrastructure | Mines |
|  |  |  | Browning |  | River fragmentation |
| *Desert*  *Semi-arid* | Browning | Human Density | Future Climate Warming |  |  |
|  |  | Livestock Density | Linear Infrastructure |  |  |
|  |  | Urbanization | Future rainfall anomaly |  |  |
|  |  |  | Greening |  |  |
|  |  |  | Agricultural Expansion |  |  |
|  |  |  | Mines |  |  |
|  |  |  | River fragmentation |  |  |
|  |  |  | Greening |  |  |
|  | Urbanization | Future rainfall anomaly | Future Climate Warming |  |  |
|  |  | Browning | Linear Infrastructure |  |  |
|  |  | Agricultural Expansion | Mines |  |  |
|  |  | Livestock Density | River fragmentation |  |  |
|  |  |  | Greening |  |  |
|  |  |  | Human Density |  |  |
| *Western Ghats* | Future Climate Warming | Agricultural Expansion | Urbanization | Browning | Linear Infrastructure |
|  | Livestock Density |  | Human Density | Greening | Mines |
|  |  |  |  | Future rainfall anomaly | River fragmentation |
| *Deccan peninsula* | Livestock Density | Urbanization | Future rainfall anomaly | Human Density | Linear Infrastructure |
|  |  | Agricultural Expansion | Future Climate Warming | Browning | Mines |
|  |  |  |  | Greening | River fragmentation |
| *Gangetic plains* | Agricultural Expansion | Future rainfall anomaly | Linear Infrastructure | Human Density |  |
|  | Future Climate Warming | Urbanization | Livestock Density |  |  |
|  |  | Greening | Mines |  |  |
|  |  | Browning | River fragmentation |  |  |
| *Northeast* | Livestock Density | Greening | Future Climate Warming | Linear Infrastructure | Browning |
|  |  |  | Urbanization | Human Density | Future rainfall anomaly |
|  |  |  | Agricultural Expansion | Mines |  |
|  |  |  |  | River fragmentation |  |

Notes: We started by collectively identifying the entire suite of threats that could impact biodiversity, habitats and ecosystem services. We then streamlined these to a set of 11 threats that encompassed: direct impacts of human presence, land-use change (e.g., urbanization and agricultural expansion), high-intensity disturbances such as mines and linear infrastructure, and climate change. Our next consideration was the availability of spatially-explicit data on these threats for all the focal regions of this study. We sourced country-wide spatial layers for each of these attributes (see Table S2). For each zone, all assessors (authors of the current study) ranked the threats in order of impact for the corresponding zone. Authors restricted their input to only those zones where they had field experience and expertise. We then averaged ranks across all assessments. We handled ties and close rankings by grouping them together into single ranks. Finally, we reviewed these ranks zone-wise, collectively, to arrive at the final consensus ranking used in the above table.
